# FEASIBILITY OF VASCULAR REMODELING PARAMETER ESTIMATION FOR ASSESSING HYPERTENSIVE PREGNANCY DISORDERS

**DOI:** 10.1101/2021.11.10.468132

**Authors:** Georgios Kissas, Eileen Hwuang, Elizabeth W. Thompson, Nadav Schwartz, John A. Detre, Walter R. Witschey, Paris Perdikaris

**Affiliations:** Department of Mechanical Engineering and Applied Mechanics, University of Pennsylvania, Philadelphia, PA, USA; Department of Bioengineering, University of Pennsylvania, Philadelphia, PA, USA; Maternal Fetal Medicine, Department of Obstetrics and Gynecology, Perelman School of Medicine, University of Pennsylvania, Philadelphia, PA, USA; Department of Radiology, Perelman School of Medicine, University of Pennsylvania, Philadelphia, PA, USA; Department of Neurology, Perelman School of Medicine, University of Pennsylvania, Philadelphia, PA, USA

**Keywords:** Bayesian inference, 4D Flow MRI, Reduced order modeling, Hypertension, Pregnancy

## Abstract

Hypertensive pregnancy disorders, such as preeclampsia, are leading sources of both maternal and fetal morbidity in pregnancy. Non-invasive imaging, such as ultrasound and magnetic resonance imaging (MRI), is an important tool in predicting and monitoring these high risk pregnancies. While imaging can measure hemodynamic parameters, such as uterine artery pulsatility and resistivity indices, the interpretation of such metrics for disease assessment rely on ad-hoc standards, which provide limited insight to the physical mechanisms underlying the emergence of hypertensive pregnancy disorders. To provide meaningful interpretation of measured hemodynamic data in patients, advances in computational fluid dynamics can be brought to bear. In this work, we develop a patient-specific computational framework that combines Bayesian inference with a reduced-order fluid dynamics model to infer remodeling parameters, such as vascular resistance, compliance and vessel cross-sectional area, known to be related to the development of hypertension. The proposed framework enables the prediction of hemodynamic quantities of interest, such as pressure and velocity, directly from sparse and noisy MRI measurements. We illustrate the effectiveness of this approach in two systemic arterial network geometries: an aorta with carotid and a maternal pelvic arterial network. For both cases, the model can reconstruct the provided measurements and infer parameters of interest. In the case of the maternal pelvic arteries, the model can make a distinction between the pregnancies destined to develop hypertension and those that remain normotensive, expressed through the value range of the predicted absolute pressure.

## 1 Introduction

Hypertensive pregnancy disorders (HPDs) increase risk to health in mothers and infants. It is estimated that 5% of pregnancies in the United States are complicated by preeclampsia, the most common form of HPD, and rates have been steadily increasing per year^1^. However, the pathophysiology of HPD is poorly understood, leaving clinicians without a screening test to reliably assess risk of adverse pregnancy outcomes^2^. While several studies have investigated the utility of biochemical markers and/or ultrasound (US) in early prediction of HPD, their positive predictive value has not been high enough for standard clinical practice^3^. Even if HPD could be accurately predicted, there are still limited treatment options for HPD due to under-investigation of the nature of the disorder^4^.

It has been suggested that insufficient spiral artery remodeling during placental development may be the cause of high blood pressure and other hemodynamic disturbances in HPD^5^. Since the latest non-invasive in vivo imaging technologies have not been able to resolve the small spiral arteries (diameters on the order of ~ 10^2^ *μm*, which is currently smaller than the resolution of MRI for body imaging), a popular alternative research tool has been Doppler US velocimetry of the uterine arteries (UtAs), which are upstream from the spiral arteries and easier to visualize because of their larger diameter^6^. More recently, 4D flow MRI velocimetry of the UtAs has been investigated^7,8^. Compared to US, 4D flow MRI has lower temporal resolution and longer scan time but larger spatial coverage, improved spatial resolution, and measures velocity in three dimensions rather than only one.

In normal pregnancy, the UtAs remodel outwardly, with larger vessel lumen area and little to no change in wall thickness^9,10^. This is mediated by a combination of local wall shear stress, nitric oxide release, and local/systemic endocrine signaling. Animal studies on tissue reorganization within the UtA wall have observed hyperplasia and hypertrophy of the smooth muscle cells to maintain wall thickness during vasodilation^10,11^. Changes in elastin and collagen content in the UtA wall have been found to be variable in different animal species^10^. One study reported evidence of increased myogenic tone in human pregnancy, which is surprising given the amount of vasodilator molecules released^12^. Nevertheless, this appears to signify that the uteroplacental arteries play a critical role in regulating the distribution of blood flow during pregnancy.

Doppler US velocimetry has shown that UtA flow increases with gestational age while pulsatility and resistivity indices (PI and RI) decrease with gestational age in normal pregnancy^6,13^. The PI corresponds to the difference between the maximum and minimum blood velocity, normalized by the mean velocity and the RI corresponds to the difference between the maximum and the minimum blood velocity normalized by the maximum blood velocity. In early pregnancy, the velocity waveform may contain a diastolic notch, representing the reflected blood flow of uteroplacental circulation^14^, which is expected to diminish by third trimester as the uteroplacental vasculature continues to remodel^13^. These features suggest that persistently high pulsatility and the presence of the diastolic notch would be associated with adverse pregnancy outcomes. However, despite numerous investigations, US measurement of UtA velocity still has low positive predictive value, which has precluded it from entering routine clinical practice^3^.

Computational fluid dynamics (CFD) has been a useful research tool to help identify sources of UtA velocity waveform changes and perhaps shed light on any biases that may weaken its diagnostic and screening utility. The earliest models were 0D lumped parameter models based on Frank’s Windkessel model, which allowed mathematical descriptions of hemodynamic parameters using physical principles adopted from electrical circuits^15^. Fundamentally, the Windkessel model relates pressure (*P*) to flow (*Q*) as *P* = *QR*, where *R* denotes resistance. 1D models have also been found to be computationally efficient and validated for accuracy in various arterial networks and flow conditions^16,17^. More recently, advances in computing have enabled 3D fluid-structure simulations of blood flow in localized organs and tissues^18,19,20^.

Simulations of UtA flow reported that high resistance and small UtA radius recapitulated the abnormal waveform shapes (high PI with diastolic notch) that closely matched measurements acquired from a preeclamptic mother^21,22^. CFD methods were also leveraged to investigate spiral artery remodeling as a potential source of the abnormal waveform shapes detected in UtA US. While insights were gained in the interaction between trophoblast invasion and hemodynamic changes of spiral arteries during pregnancy^5,23^, evidence suggests that PI and the diastolic notch may be a moderate proxy for trophoblast invasion but other factors such as radial artery and arteriovenous anastomoses remodeling may also contribute to the waveform shape^24^. This suggests the need for more investigation into the complex remodeling-hemodynamic interactions underlying HPD that would lead to more effective clinical biomarkers than PI/RI. Previous uterine artery computational modeling has sought to understand the physical mechanisms underpinning measured ultrasound velocity wave-forms^25^, but without considering a patient specific large vessel structure in the female pelvis.

In this study, we leverage a hybrid 0D-1D model first proposed by *Sherwin et. al.*^26^ to derive remodeling parameters, such as arterial resistance, compliance, and cross-sectional area, from non-invasive imaging flow measurements of the proximal uterine arteries for patient specific pelvic geometries. To achieve this, we developed a computational framework that combines a reduced order Navier-Stokes model as a forward evaluation model and a Markov Chain Monte Carlo (MCMC) algorithm for predicting these biological indices that could be later used to assess the progression of HPD. We first demonstrate feasibility in the aorta with synthetic and measured MRI flow data, followed by a similar analysis of the maternal pelvic arterial network of a normal and a pregnant subject that went on to develop hypertension later in pregnancy after MRI. We also compare the corresponding PI and RI from MRI and US in these subjects.

In a Bayesian estimation setting, reduced order models are commonly coupled with MCMC algorithms for inferring posterior distributions over the unknown parameters given noisy measurements. For example, Larson *et. al.*^27^ introduced a Bayesian model selection algorithm for identifying the position of wall abnormalities in a complex arterial network. Their algorithm employed a transition MCMC strategy coupled with a reduced order flow model in order to examine different flow model setups, chose the one that best described the underlining phenomena, and matched the target waveform. Thus, the objective of this work was to solve a classification problem, where the boundary conditions of the problem were assumed and the geometrical and structural properties of the vessel were varying. Moreover, Colebank *et. al.*^28^ proposed a strategy that employed a Delayed Rejection Adaptive Metropolis (DRAM) algorithm in order to infer the boundary conditions for a three-element Windkessel model in pulmonary hypertension and performed a sensitivity analysis to identify the model parameter influence on the predicted pressure. In this case, a parameter reduction strategy was employed and nominal parameter values for *R* and *C* were assumed, which then reformed the inverse problem to identifying scaling multipliers for the nominal parameters at each outlet. The authors employed pressure data for the main pulmonary artery (MPA) in order to infer the arterial resistance and compliance for the outflow vessels and performed model sensitivity analysis to assess the parameters’ importance to the model output. In another approach, Puaun *et. al.*^29^ proposed an MCMC method for inferring flow parameters in a pulmonary circulation vascular network in mice. The strategy was based on DRAM, with an additional parameter scaling technique. Moreover, they proposed starting the algorithm using a Maximum Likelihood approach in order to speed up the convergence of the algorithm and presented convergence tests in order to assess the effectiveness of the algorithm. In this work, we perform direct inference of the model parameters only via sparse, cross-sectional averages of 4D flow MRI measurements. This increases the complexity of the inference, as there is an inherent uncertainty caused by the discrepancy between the chosen model and the real flow conditions. Because of this discrepancy, different combinations of inputs and their corresponding model discrepancy might provide results that are in close agreement and thus provide high values of the likelihood function during the Monte Carlo sampling.

In summary, the main contribution of this work is a patient specific methodology for directly inferring the values of arterial resistance, compliance and equilibrium cross-sectional area, as well as estimating of the absolute pressure in maternal pelvic arteries. This is a first step towards investigating the relationship between vascular remodeling and pregnancy outcomes for early prediction of HPD.

## 2 Methods

### 2.1 MRI data acquisition and processing

#### MRI of aorta and carotid in non-pregnant volunteer

For this study, we use the same aorta data-set reported in Kissas *et al.*^30^. The MRI data was acquired from a 27-year-old healthy female volunteer using a 1.5T MRI scanner (Avanto, Siemens Healthineers, Erlangen, Germany). The protocol included a balanced steady-state free precession (bSSFP) localizer, prospectively electrocardiogram (ECG)-gated 2D cine and 2D phase contrast images prescribed at four locations in the aorta and one location in the left common carotid artery. The temporal resolution was 29.4-34.8 ms and 20.7 ms for the 2D cine and 2D phase contrast MRI, respectively. The time-resolved cross-sectional areas were extracted from the 2D cine images and time-resolved velocity wave-forms were extracted by processing the 2D phase contrast phase difference images (ImageJ v1.48; imagej.nih.gov/ij). The arc lengths between points were computed after segmentation and center-line extraction of the bSSFP localizer images using Seg3D 2.3.0 (http://sci.utah.edu/cibc-software/seg3d.html)^31^ and VMTK (http://www.vmtk.org/)^32^. Figure 1 shows the resulting geometry. Additional details about the MRI acquisition and post-processing have been reported previously^30^.

**Figure 1:**
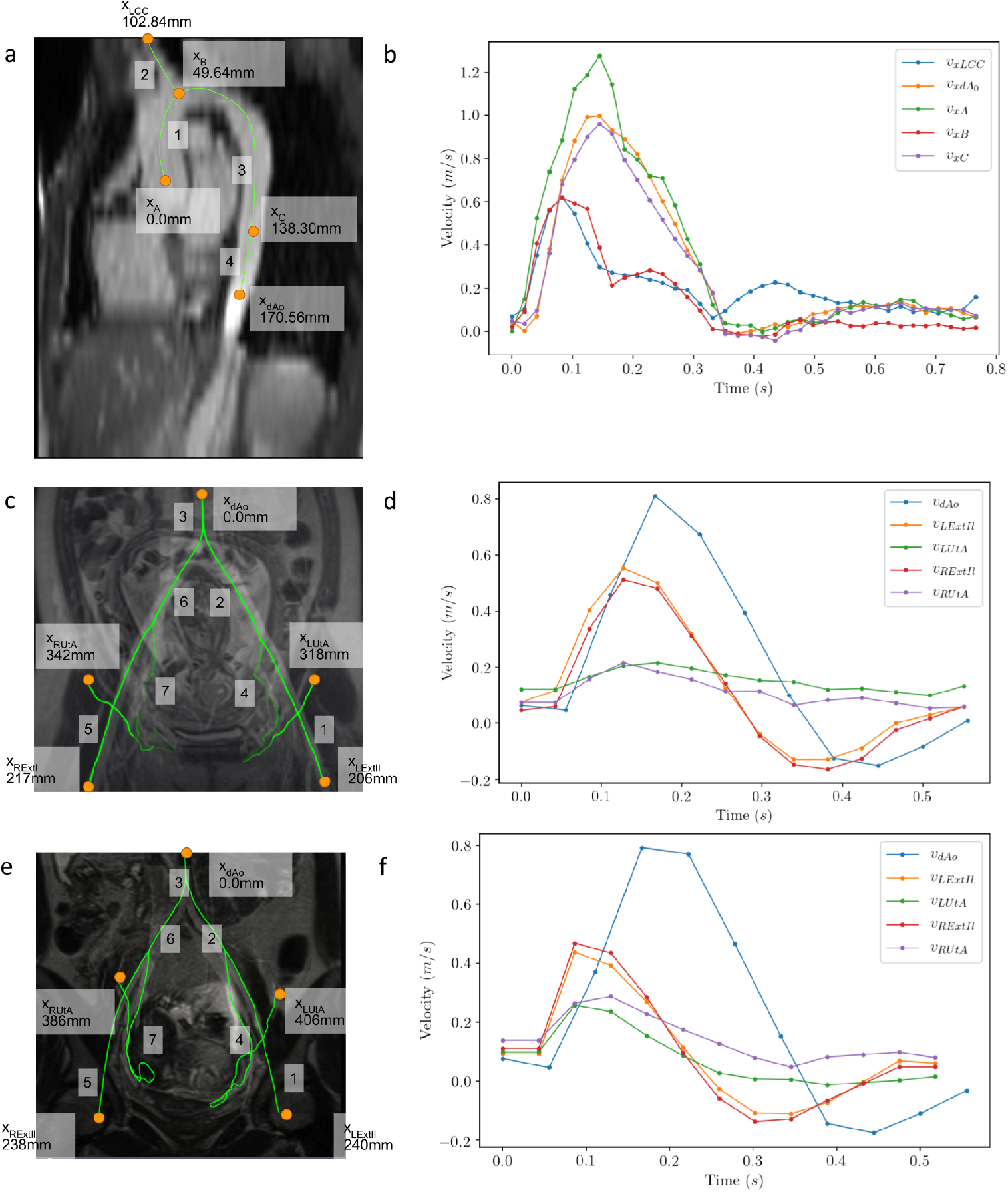
Schematic representation of the MRI geometry and velocity data of human volunteers. a) Center-line geometry with velocity measurements at points *x*_*A*_ (ascending aorta), *x*_*B*_ (aortic arch), *x*_*C*_, *x*_*dAo*_ (descending aorta outlet), and *x*_*LCC*_ (left common carotid artery outlet). Vascular segments are labeled from 1 to 4, each connecting a pair of points as shown. All lengths are reported relative to *x*_*A*_. b) Aorta velocity wave-forms corresponding to orange points in panel (a) measured from 2D phase contrast MRI. In c) the geometry of the Normal subject maternal pelvic arteries is presented and in d) the corresponding inlet and the outlet (simulation target) waveform for the orange points in panel (c) are presented. In e) the geometry of the pre-HPD maternal pelvic arteries is presented and in f) the inlet and the outlet (simulation target) wave-forms corresponding to the orange points in panel (e) are presented. For both cases of maternal pelvic geometries vessel 3 corresponds to the aorta, vessel 6 to the right common iliac artery, vessel 2 to the left common iliac artery, vessel 7 to the right uterine artery, vessel 4 to the left uterine artery, vessel 1 to the left external iliac artery and vessel 5 to the right external iliac artery.

#### MRI of uterine artery in human pregnancy

Using the same scanner, MRI data was acquired from a healthy pregnant volunteer, denoted as the Normal subject (maternal age=24 years, gestational age (GA)=18.7 weeks) and a pregnant volunteer that developed hypertension in late pregnancy, denoted as the pre-HPD subject (maternal age=25 years, GA=19.1 weeks). The pre-HPD subject was not diagnosed with chronic hypertension nor was on any related medication and was diagnosed with preeclampsia four months after the MRI data were collected. Each subject was positioned in feet-first supine with a 12-channel spine array coil and two 4-channel body matrix coils. Both patients were imaged in supine position with no tilt. An ECG monitor was attached to the subject for synchronization of the MRI to heart rate. The protocol consisted of a half-Fourier acquisition stimulated echo (HASTE) localizer of the abdomen and pelvis, a time-of-flight (TOF) angiogram of the abdomen and pelvis, a 2D prospectively-gated phase contrast scan of the descending aorta, and prospectively-gated 4D flow MRI of the uterine arteries (UtAs) and external iliac arteries (ExtIls). The details of the MRI acquisition parameters are listed in Table 1.

**Table 1:**
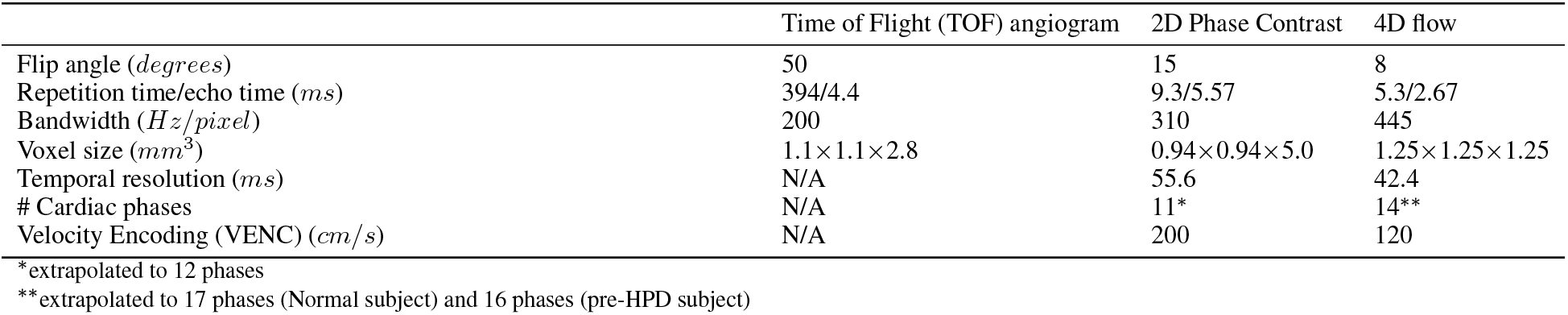
MRI parameters used to acquire geometry and velocity data for the two pregnant subjects, Normal and pre-HPD.

|  | Time of Flight (TOF) angiogram | 2D Phase Contrast | 4D flow |
| --- | --- | --- | --- |
| Flip angle (*degrees*) | 50 | 15 | 8 |
| Repetition time/echo time (*ms*) | 394/4.4 | 9.3/5.57 | 5.3/2.67 |
| Bandwidth (*Hz/pixel*) | 200 | 310 | 445 |
| Voxel size (*mm<sup>3</sup>*) | $1.1 \times 1.1 \times 2.8$ | $0.94 \times 0.94 \times 5.0$ | $1.25 \times 1.25 \times 1.25$ |
| Temporal resolution (*ms*) | N/A | 55.6 | 42.4 |
| # Cardiac phases | N/A | 11* | 14** |
| Velocity Encoding (VENC) (*cm/s*) | N/A | 200 | 120 |

\*extrapolated to 12 phases
\*\*extrapolated to 17 phases (Normal subject) and 16 phases (pre-HPD subject)

During post-processing of the MRI data, the TOF angiograms were segmented (Seg3D^31^) and center-lines were extracted (VMTK^32^) to compute path length from the descending aorta to the UtAs and ExtIls (Figure 1). The equilibrium areas of each vascular segment were estimated from multi-planar re-formats of the TOF images. Velocity and area wave-forms were extracted from 2D phase contrast images using ImageJ. The 4D flow images were processed with custom software (MATLAB,Natick,Massachusetts)^33^, followed by velocity-based thresholding of the volumetric iso-surface (Ensight, CEI; Apex, NC). Velocity wave-forms were extracted from the left/right UtAs and left/right ExtIls. Since both the 2D phase contrast MRI and 4D flow MRI were prospectively-gated, a few end-diastolic cardiac phases were not acquired from each cardiac cycle. These end-diastolic velocity values were extrapolated by averaging the velocities of the first and last cardiac phase (Table 1).

Each subject was positioned in semi-recumbent supine during ultrasound (US) scanning by a clinician (N.S.) with extensive experience in prenatal US. The UtAs were scanned using the C4-8 transabdominal probe of a GE Voluson E10 (GE Healthcare, Wisconsin, United States) US machine. Measurements of UtA Doppler velocity wave-forms were recorded bilaterally, and PI and RI were calculated for comparison with the MCMC estimation.

### 2.2 A hybrid 0D-1D pulsatile flow model

A reduced order Navier-Stokes flow model is employed to simulate pulsatile flow in a network of compliant vessels. In order to perform such reduction, a series of assumptions are made. Blood is approximated as a Newtonian, incompressible fluid, with constant viscosity and density. The vessels are approximated as thin, impermeable, elastic, axi-symmetric compliant cylinders with much larger length than the radius (*L/r* ≫ 1), which is restricted to radial displacement. The pressure and structural properties of the vessel do not vary across its cross-section^34,26^. By averaging the conservation of mass and momentum over the cross-sections, a system of hyperbolic conservation laws is derived^26^. Considering the vessel to be a thin-walled cylinder, Laplace’s law is used to derive an equation that relates the pressure with the structural properties of the vessel^34^. This equation is used to calculate the absolute pressure value. The resulting system of equations takes the form

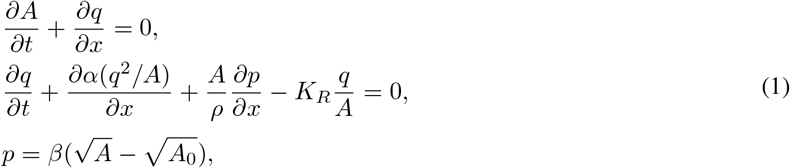

where *p*(*x, t*), *q*(*x, t*) and *A*(*x, t*) represents the pressure, the flow and cross-sectional area, respectively. Moreover, 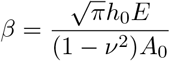 and *A*_0_ denotes the vessel’s cross-sectional area at equilibrium, *E* the Young’s modulus, *h*_0_ the wall thickness, *ν* the Poisson ratio and *ρ* the blood density. Values for *h*_0_, *E* and *ν* can be found in the literature, while *A*_0_ is measured via MRI. Moreover, *K*_*R*_ is a friction parameter which depended on the chosen velocity profile and *α* is a momentum flux correction parameter which accounts for the non-linearity of the sectional integration in terms of the local velocity^35^. In this study, *K*_*R*_ = −22*μπ*, where *μ* is the dynamic blood viscosity and *α* = 1.1. For all experiments, *β* is computed based on the empirical relation reported by Olufsen *et. al.*^36^. The tortuosity of the geometry is assumed to be small enough such that the system (1) is valid, but in reality there was some uncertainty induced by neglecting the complex topology, particularly of the uterine arteries.

In a network of arterial vessels, a series of domain conditions regarding the relationship between vessel segments need to be satisfied. A pulsatile flow wave coming from the heart is set as the inflow boundary condition. In this study, the pulsatile inflow wave is defined by MRI data, which then smoothed using Gaussian Process regression with a periodic kernel^37^, and subsequently approximated using a Fourier series expansion. At locations of vessel bifurcation, it is assumed that there existed no mass leakage so the conservation of mass between the parent (i.e. vessel # 1) and the daughter vessels (i.e. vessels #2, #3) has to be satisfied. The mass conservation equation is defined as:

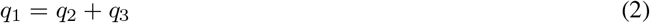

where *q*_1_ and *q*_2_, *q*_3_ are the flow in the parent and daughter vessels, respectively. The bifurcation is considered to occur at a point and by assuming laminar flow, no energy loss, and no gravity effects, Bernoulli’s law is employed for the inlet and outlets of the arterial network. It is shown that the pressure continuity *p*_1_ = *p*_2_ = *p*_3_ is a sufficient condition to produce accurate results for geometries consisting of large vessels^34^, so by combining this assumption with Bernoulli’s law, the following relation is obtained:

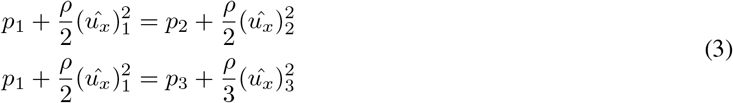

where 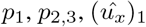, and 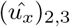 are the pressure and velocity values of the parent and daughter vessels, respectively. This constitutes the full forward model for predicting the target quantities (flow, wall displacement, and absolute pressure). We employ an in-house Discontinuous Galerkin (DG) solver^26,38^ to compute the solution of this model. We extract the vessel centerlines from the three-dimensional arterial geometry of each patient and created a graph as the input geometry of the DG simulator.

For the DG simulation, detailed boundary conditions are necessary to accurately describe the underlying physical phenomena. In this study, a three-element Windkessel model^39^ is employed having a governing equation:

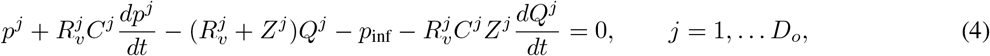

where *D*_*o*_ represents the total number of outlets, *Q*^*j*^ = *A*^*j*^ *u*^*j*^ is the flow rate at the outlet, and *p*_inf_ = 666.5 Pa denotes the constant downstream pressure. *Z*^*j*^ represents the characteristic impedance^40^ that is defined as 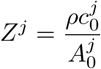 where *C*^*j*^ is the total arterial compliance and 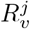 is the systemic vascular resistance of vessel #j. The characteristic impedance is chosen in such a way that allows the incoming wave to reach 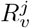 and *C*^*j*^ without being reflected^40^. For modeling the circulation of an arterial network the most important challenge is related to correctly prescribing the 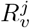 and *C*^*j*^ parameters, such that the resulting predictions are within a physiologically accurate range. In this study, we are estimating the total arterial resistance (*R*^*j*^), defined as:

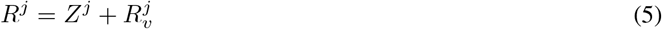

for each vessel *j*.

### 2.3 Bayesian Inference

The proposed pipeline for statistically estimating unknown parameters for each outlet using MRI measured velocity data consists of two building blocks, the forward model and the sampling method. The forward model is the hybrid 0D-1D blood flow solver discussed in section (2.2). Its input is a set of parameters *θ*^*j*^ determined by the sampling method and the output is a prediction of the quantities of interest to be compared with the observed clinical data (in our case the velocity). In this work, we employ slice sampling^41^, because it is a technique that can be used as a “black-box” sampler, meaning that requires little tuning, compared to Metropolis based methods, can effectively adjust the step size to match the local shape of the density function and can be as efficient as a Gibbs sampler without having to derive the conditional distributions. A comprehensive overview of the method, as well as, a comparison with existing popular sampling methods can be found in the original work of Neal *et. al.*^41^. The model parameters are defined as a triplet 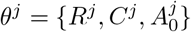 for each outlet #*j*. Under this setting, the target posterior distribution is expressed as:

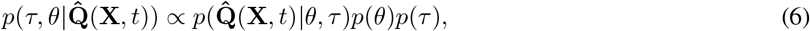

where 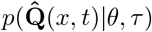 denotes the likelihood function and *τ* the measurement noise variance. *p*(*θ*) and *p*(*τ*) denote the prior distributions over the unknown parameters and the measurement noise variance, respectively. Assuming independence across the *N*_*o*_ observed measurements and a Gaussian noise model, the likelihood is factorized as:

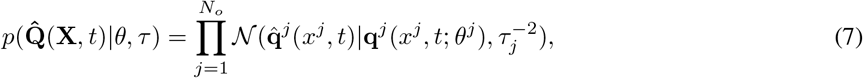

where *N*_*o*_ is the number of outlets, 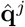 represents the target measurements, **q**^*j*^ represents the forward model prediction for parameter set *θ*^*j*^, and *τ*_*j*_ represents the noise variance of the measurements for vessel #*j*. The use of the Gaussian likelihood is based on the assumption that the measurements may be corrupted by Gaussian noise with zero mean and variance 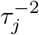.

In the Bayesian inference algorithm, we chose the prior parameter distributions based on our prior knowledge of how the parameters are distributed. The prior distributions for the model parameters were defined as:

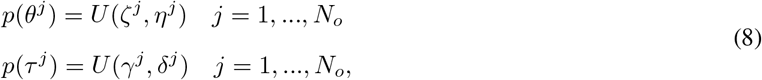

where *ζ*^*j*^, *η*^*j*^ are user-defined lower and upper bounds, respectively, of the considered parameter range of the considered outlet. Furthermore, *γ*^*j*^ is the lower and *δ*^*j*^ the upper bounds for the noise parameter for each outlet. The uniform distribution is chosen for the priors^42^ to limiting the chance in which the distribution mass may be concentrated to a particular region. In the case that information over the prior distribution form and range is known, informative priors can be employed in order to accelerate the sampling convergence.

The Bayesian inference algorithm discovers sets of parameters lying in regions of high probability under the posterior distribution 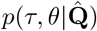 By approximating this distribution we are able to make predictions about the quantities of interest and at the same time quantify the predictive uncertainty of our parameter estimates. In order to produce realizations of the system outputs we define:

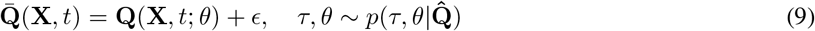

where 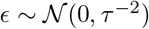 the inferred noise distribution. We can, also, define the maximum-a-posteriori (MAP) probability estimate and quantify the corresponding uncertainty of the model by sampling parameters out of the posterior distribution. By sampling the posterior distribution we can then estimate the mean and the variance of the prediction in order to assess its uncertainty. The mean is defined as

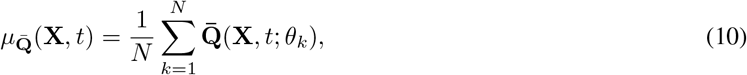

while the variance is defined as

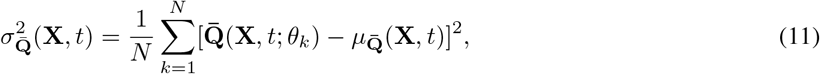

where *N* denotes the number of posterior samples. The above algorithm provides a statistical description of the predictions, which later is used to assess the quality of our conclusions.

Flow rates are computed by multiplying the predicted velocity wave-forms with the predicted cross-sectional area for each vessel. Then, they are compared to the target flow rates, which are computed by multiplying the cross-sectional area measured from TOF MRI with the velocity measured from 4D flow MRI. The algorithm is schematically presented in figure 2.

**Figure 2:**
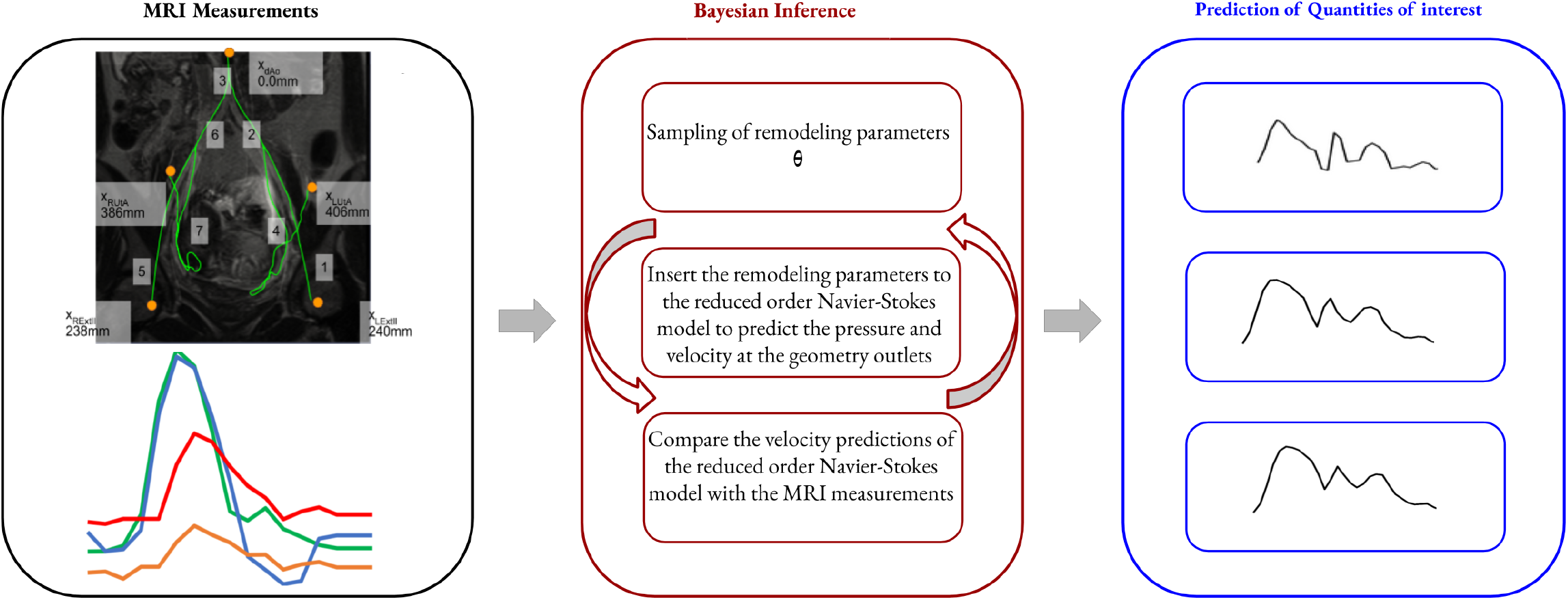
Schematic representation of the process proposed in the manuscript. The procedure starts by considering MRI velocity measurements and information about the geometry of a patient, then samples remodeling parameters based on a sampling algorithm, which it uses to make velocity predictions followed by comparison with the input measurements. Upon convergence to a stationary distribution the parameters are used in order to make velocity and pressure predictions.

### 2.4 Experimental Setup

The Bayesian inference algorithm was tested on three MRI volunteer data-sets, one collected in the aorta and two in the maternal pelvic arteries (Section 2.1). For all experiments, predefined parameters were *K*_*R*_ = −22*μπ* and *α* = 1.1 to account for viscous losses, blood density *ρ* = 1060 Kg/m^3^, and viscosity *μ* = 3.5 mPa s. The code for all the methods is created by combining the python library for Bayesian statistical modeling PyMC3^43^ with a custom C language code including a custom loss function in the python library Theano^44^.

#### 2.4.1 Aorta Experiments

The arterial network geometry consists of four vessel segments with one inlet (ascending aorta), a bifurcation, and two outlets (left common carotid artery (LCC) and descending aorta (dAo)), as shown in Figure (1). Given this data-set we perform two computational experiments, one using synthetic velocity data and one using MRI measured velocity data. This aorta data-set was originally reported in a previous study^30^.

In the first aorta experiment, the geometry was coupled with synthetic velocity data at the outlets to formulate a simplified problem as a proof-of-concept of the proposed Bayesian inference framework. To construct the synthetic data, the Discontinuous Galerkin (DG) simulator was provided with an arbitrary set of Windkessel parameters (*R*, *C*). The generated velocity data was then corrupted with 2% Gaussian noise to simulate MRI measurement error. Despite neither the noise level being that small nor the noise model Gaussian in real cases, we consider this case as a sanity check, thus we define it in a simple manner. Moreover, we consider the model discrepancy^45^, the difference in approximation quality between the employed and the real physical models that describe the flow, as the main factor contributing to the prediction uncertainty. The period of the volunteer’s cardiac cycle *T* = 0.78 s was preserved in the simulation but *N*_*p*_ = 50 points were arbitrarily chosen to represent the waveform. Further details of the simulation setup have been previously discussed in^30^.

The Bayesian inference algorithm is set up to take the geometry and synthetic data and estimate the Windkessel parameters and pressure wave-forms. The search space is defined by uniform prior distributions for the unknown parameters (see Table 2) chosen based on the intuition described in Section 2. The area wave-forms provided by MRI are not taken into consideration for simplicity of this proof-of-concept experiment, but MRI measurements of equilibrium cross-sectional area are used to determine the wall stiffness parameter *β* in the one-dimensional blood flow model. To assess accuracy after convergence, the estimated parameters are compared with the target parameters *R*_*dAo*_ = 1.667 × 10^8^ *Pa s m*^−3^, *C*_*dAo*_ = 9.002 × 10^−9^ *Pa*^−1^ *m*^3^, *R*_*LCC*_ = 2.102 × 10^9^ *Pa s m*^−3^, and *C*_*LCC*_ = 2.538 × 10^−10^ *Pa*^−1^ *m*^3^, which are taken based on the findings reported in^30^.

**Table 2:**
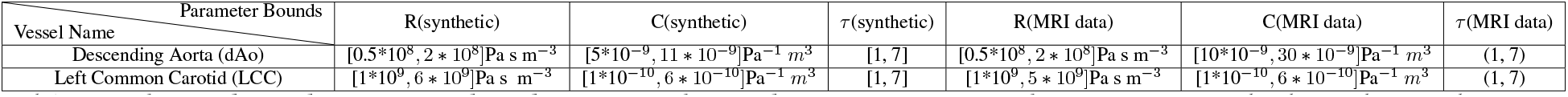
Chosen bounds for prior distributions of the model parameters for the aorta case: The bounds are chosen as described in section 2, for the total arterial resistance (R) and compliance (C). Initial knowledge from^30^ was incorporated into the setup of these ranges.

In the second aorta experiment, the same geometry is coupled with the measured MRI velocity data at the outlets in order to estimate the set of Windkessel parameters that provide the best match for the given target measurements. Prior to starting the Bayesian inference algorithm, a Gaussian process regression^37^ with a periodic kernel is performed on the MRI wave-forms to ensure periodicity, to augment the *N*_*p*_ = 38 points to *N*_*p*_ = 100 to improve precision and to align the predicted velocity and MRI velocity wave-forms which have different temporal resolutions and starting times. More details about the pre-processing of the MRI data are described in^30^. The Bayesian inference algorithm is set up with a search space defined by the uniform prior distributions reported in Table 2.

#### 2.4.2 Maternal pelvic arterial experiments

The Bayesian inference methodology is performed on two data-sets comprised of MRI measurements of velocity from the female pelvic region of two human volunteers. The first data-set corresponds to the Normal subject and second to the pre-HPD subject, see Section 2. The maternal pelvic arterial network geometry consists of seven vessels extending from the descending aorta (dAo) to the right and left common iliac arteries (RCI, LCI), to the right and left uterine arteries (RUtA, LUtA) and external iliac arteries (RExtIl, LExtIl). Each network contained one inlet, four outlets, and two bifurcations, as shown in Figure (1).

For each network, the Bayesian inference algorithm is provided with MRI measured velocities at each outlet and then executed to estimate the Windkessel parameters *R* and *C* as well as the equilibrium cross-sectional area (*A*_0_), similar to the aorta experiments. Unlike the aorta experiments, cross-sectional area measurements are included in the inference. In contrast to the descending aorta region where the cross-sectional area magnitude of the vessels is large enough to be accurately captured by the MRI resolution, in the pelvic region the arteries are very narrow and thus inaccuracies are introduced in the measurements. In the maternal pelvic experiments, the structural MRI measured cross-sectional area of the vessels that do not correspond to outlets (i.e. descending aorta, left and right common iliac arteries) is inserted to the model and kept constant throughout the entire inference procedure, because we do not have enough data to infer them. Therefore, we consider the cross-sectional area as a free parameter for the outlets to potentially correct for the induced inaccuracies in the geometry of their parent vessels.

Since 4D flow MRI consists of velocity measurements from planes along the vessel cross section at different locations, we can construct time series for both the maximum and the mean velocity. In standard practice, the mean velocity wave-forms are used to compute the mean flow rate. However, the mean velocity wave-forms have low pulsatility therefore the maximum velocity is used in practice for calculating Pulsatility Index (PI) and Resistivity Index (RI). The reduced order model employed in this work is a cross-sectionally averaged version of the Navier-Stokes equation, so we will use the mean MRI velocity for estimating the pressure wave-forms, *R*, *C*, and *A*_0_. Moreover, we will run a separate Bayesian inference experiment for the sole purpose of calculating Pulsatility Index (PI) and Resistivity Index (RI) to compare with US measurements. This computation constitutes another sanity check for assessing the quality of the approximation that the framework can achieve for the case of predicting biomarkers. The biomarkers are computed using the mean, the minimum and the maximum of the velocity waveform, therefore we can assess how well the model can approximate these waveform characteristics.

For both volunteers, the measured 4D flow MRI velocity wave-forms have a low temporal resolution, they consist of only *N*_*p*_ = 14 points over a cardiac cycle of *T* = 0.612 s. To this end, the MRI velocity wave-forms are augmented by a Gaussian process with a periodic kernel regression to create a set of *N*_*p*_ = 55 points. This additional step is beneficial to the procedure because by creating a larger baseline periodic data-set we can have stronger comparison with the prediction which results to a more accurate algorithm. Moreover, this step is used to align the velocity time-series produced by the model and the MRI measurements, so that the predictions and the measurements are on the same times. Table 3 contains the bounds of the uniform prior distributions predefined for the Bayesian inference framework employed for the Normal subject and the pre-HPD subject.

**Table 3:** Chosen bounds for the prior distributions of the model parameters used for the two maternal pelvic arterial experiments: In this table, we present the parameter bounds for the total arterial resistance (*R*), total arterial compliance (*C*) and equilibrium cross-sectional area (*A*). The orders of magnitude of the total arterial resistances and compliances are chosen based on the results from^30^ and the ranges based on our experience with simulations, the ranges of the equilibrium cross-sectional areas are chosen based on the literature. The abbreviation LExtIl correspond to the Left External Iliac, RExtIl correspond to the Right External Iliac, LUtA correspond to the Left Uterine and RUtA to the Right Uterine arteries.

| Parameter Bounds<br>Vessel Name | R | C | A |
| --- | --- | --- | --- |
| LExtII (Normal subject) | $[1*10^8, 65 * 10^8] \text{ Pa s m}^{-3}$ | $[1*10^{-11}, 255 * 10^{-11}] \text{ Pa}^{-1} \text{ m}^3$ | $[1*10^{-6}, 70 * 10^{-6}] \text{ m}^2$ |
| LUtA (Normal subject) | $[1*10^8, 65 * 10^8] \text{ Pa s m}^{-3}$ | $[1*10^{-12}, 255 * 10^{-12}] \text{ Pa}^{-1} \text{ m}^3$ | $[1*10^{-6}, 70 * 10^{-6}] \text{ m}^2$ |
| RExtII (Normal subject) | $[1*10^8, 65 * 10^8] \text{ Pa s m}^{-3}$ | $[1*10^{-11}, 255 * 10^{-11}] \text{ Pa}^{-1} \text{ m}^3$ | $[1*10^{-6}, 70 * 10^{-6}] \text{ m}^2$ |
| RUtA (Normal subject) | $[1*10^8, 65 * 10^8] \text{ Pa s m}^{-3}$ | $[1*10^{-12}, 255 * 10^{-12}] \text{ Pa}^{-1} \text{ m}^3$ | $[1*10^{-6}, 70 * 10^{-6}] \text{ m}^2$ |
| LExtII (pre-HPD subject) | $[1*10^8, 115 * 10^8] \text{ Pa s m}^{-3}$ | $[1*10^{-11}, 45 * 10^{-11}] \text{ Pa}^{-1} \text{ m}^3$ | $[1*10^{-6}, 70 * 10^{-6}] \text{ m}^2$ |
| LUtA (pre-HPD subject) | $[1*10^8, 135 * 10^8] \text{ Pa s m}^{-3}$ | $[1*10^{-12}, 45 * 10^{-12}] \text{ Pa}^{-1} \text{ m}^3$ | $[1*10^{-6}, 70 * 10^{-6}] \text{ m}^2$ |
| RExtII (pre-HPD subject) | $[1*10^8, 115 * 10^8] \text{ Pa s m}^{-3}$ | $[1*10^{-11}, 45 * 10^{-11}] \text{ Pa}^{-1} \text{ m}^3$ | $[1*10^{-6}, 70 * 10^{-6}] \text{ m}^2$ |
| RUtA (pre-HPD subject) | $[1*10^8, 115 * 10^8] \text{ Pa s m}^{-3}$ | $[1*10^{-12}, 45 * 10^{-12}] \text{ Pa}^{-1} \text{ m}^3$ | $[1*10^{-6}, 70 * 10^{-6}] \text{ m}^2$ |

## 3 Results

### 3.1 Aorta results

We present the velocity and the pressure wave-forms for the case of the synthetic data in Figure 3 and the resulting discovered parameters in Table 4. In the aorta model with synthetic velocity data, the predicted outlet velocity wave-forms closely matched the target synthetic wave-forms, see Figure 3a,b. We draw 200 samples of the posterior parameter distribution and calculated the flow rate using the predicted velocity and area. For this case, the flow rate resulting from the parameters with the maximum probability closely matched the target. For the descending aorta the model prediction is 4577.9 ± 8.1 and the target 4578.8 mL/min which results to −0.02% error. For the left common carotid artery, the predicted flow is equal to 462.8 ± 6.8 and the target 466.6 mL/min which results to −0.82% error. The predicted Windkessel parameters also closely match the target values for the left common carotid (resistance R = 2.08E+09 vs. 2.10E+09 *Pa s m*^−3^, −1.09% error; compliance C=2.54E-10 vs. 2.54E-10 *Pa*^−1^ *m*^3^, 0.12% error) and descending aorta (R = 1.67E+08 vs. 1.67E+08 *Pa s m*^−3^, 0.24% error; compliance C=8.98E-09 vs. 9.00E-09 *Pa*^−1^ *m*^3^, −0.28% error) (Table 4). In addition, the aorta model provided absolute pressure wave-forms for both outlets (see Figure 3c,d). Although validation wave-forms are not available from the measured MRI data, the predicted pressure time-series are within physiological range. The absolute systolic/diastolic pressure is 115.2 ± 1.4 / 59.7 ± 1.5 mmHg in the descending aorta and 108.0 ± 1.4/61.8 ± 1.5 mmHg for the left common carotid artery.

**Table 4:** Windkessel parameter identification for the Aorta geometry: This table reports the total arterial resistance (R) and compliance (C) discovered by the algorithm for both aorta experiment using synthetic and MRI data.The abbreviation dAo correspond to the Descending Aorta and LCC to the Left Common Carotid arteries.

| Vessel Name \ Param | $R_{\text{target}}$ | $C_{\text{target}}$ | $R_{\text{predicted}}$ | $C_{\text{predicted}}$ | Error |
| --- | --- | --- | --- | --- | --- |
| dAo(synthetic) | $1.667\text{e}+08 \text{ Pa s m}^{-3}$ | $9.002\text{e}-09 \text{ Pa}^{-1} \text{ m}^3$ | $1.671\text{e}+08 \text{ Pa s m}^{-3}$ | $8.977\text{e}-09 \text{ Pa}^{-1} \text{ m}^3$ | -1.09 % |
| LCC (synthetic) | $2.102\text{e}+09 \text{ Pa s m}^{-3}$ | $2.538\text{e}-10 \text{ Pa}^{-1} \text{ m}^3$ | $2.079\text{e}+09 \text{ Pa s m}^{-3}$ | $2.541\text{e}-10 \text{ Pa}^{-1} \text{ m}^3$ | 0.24 % |
| dAo (MRI data) | - | - | $1.150\text{e}+08 \text{ Pa s m}^{-3}$ | $12.887\text{e}-09 \text{ Pa}^{-1} \text{ m}^3$ | |
| LCC (MRI data) | - | - | $1.452 \text{ e}+09 \text{ Pa s m}^{-3}$ | $1.877 \text{ e}-10 \text{ Pa}^{-1} \text{ m}^3$ | - |

**Figure 3:**
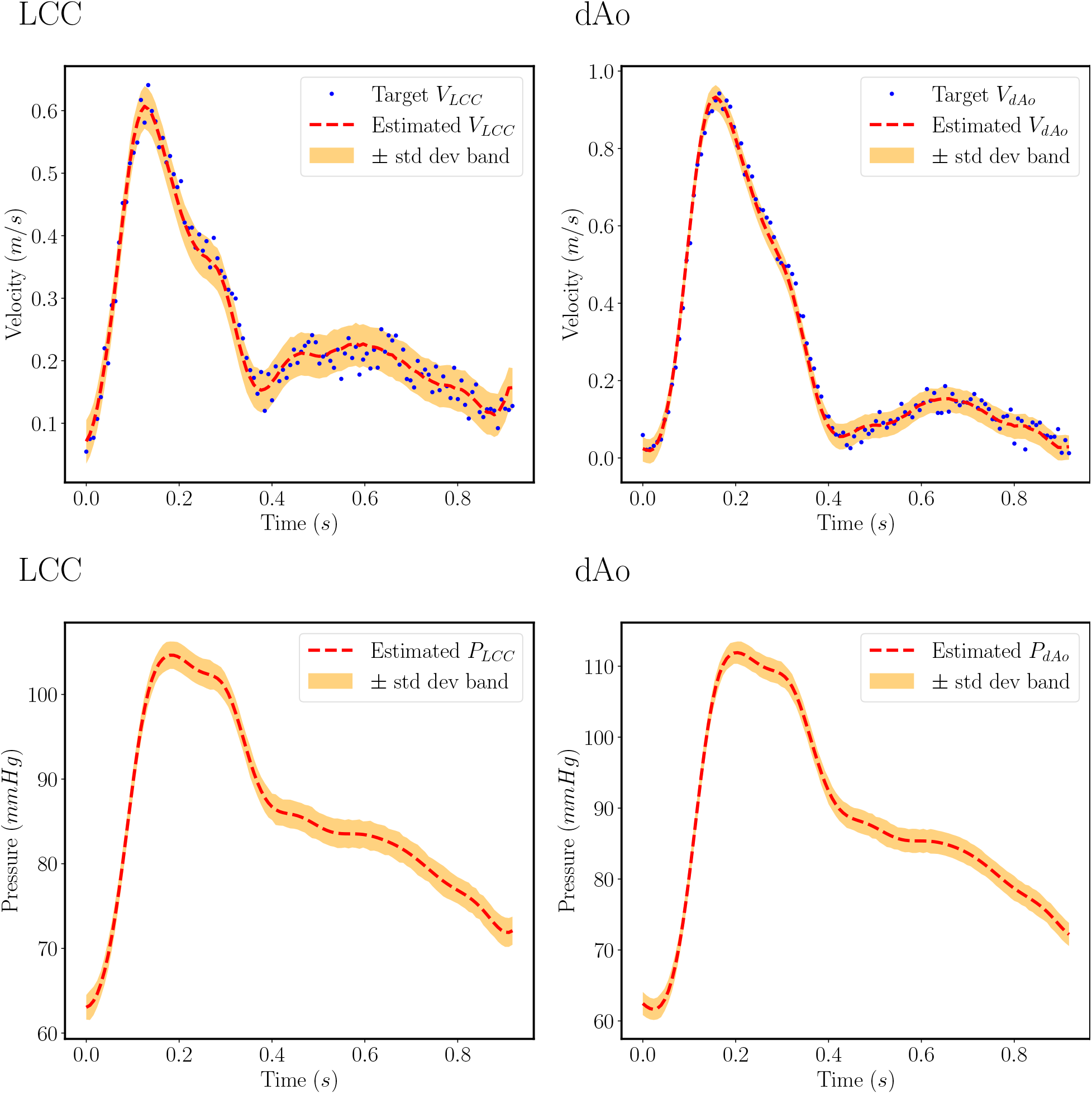
Resulting velocity and pressure wave-forms from aorta experiment using synthetic flow data: In this figure the estimated wave-forms of the aorta geometry with synthetic data is compared with the synthetic data with 2% noise corruption. The synthetic data are denoted by blue dots. The red line represents the waveform that results from using the parameters that provide the maximum likelihood and the yellow area represents the model uncertainty of the solution.The abbreviation dAo correspond to the Descending Aorta and LCC to the Left Common Carotid arteries.

When the MRI measured velocity data is used in place of synthetic data, the aorta model still achieved close agreement with the target MRI measurements. The predicted flow rate in the descending aorta is 4803.9 ± 17.5 vs. 4519.1 mL/min, 6.3 % error and in the left common carotid artery it is 353.3 ± 6.8 vs. 343.7 mL/min, 2.8% error (Figure 4a,b). Although ground truth the resistance and compliance values are not known for this particular data set, the inferred parameters can be determined the descending aorta resistance is 1.45E+09 *Pa s m*^−3^ and compliance is 1.88E-10 *Pa*^−1^ *m*^3^. The predicted resistance and compliance in the left common carotid artery are 1.15E+08 *Pa s m*^−3^ and 1.29E-08 *Pa*^−1^ *m*^3^, respectively, see Table 4. The predicted pressure wave-forms are also within physiological range^46^. The systolic/diastolic pressures in the descending aorta are 102.5 ± 1.0 / 55.1 ± 1.0 mmHg and in the left common carotid artery pressure are 95.4 ± 0.7 / 55.1 ± 0.8 mmHg (see Figure 4c,d).

**Figure 4:**
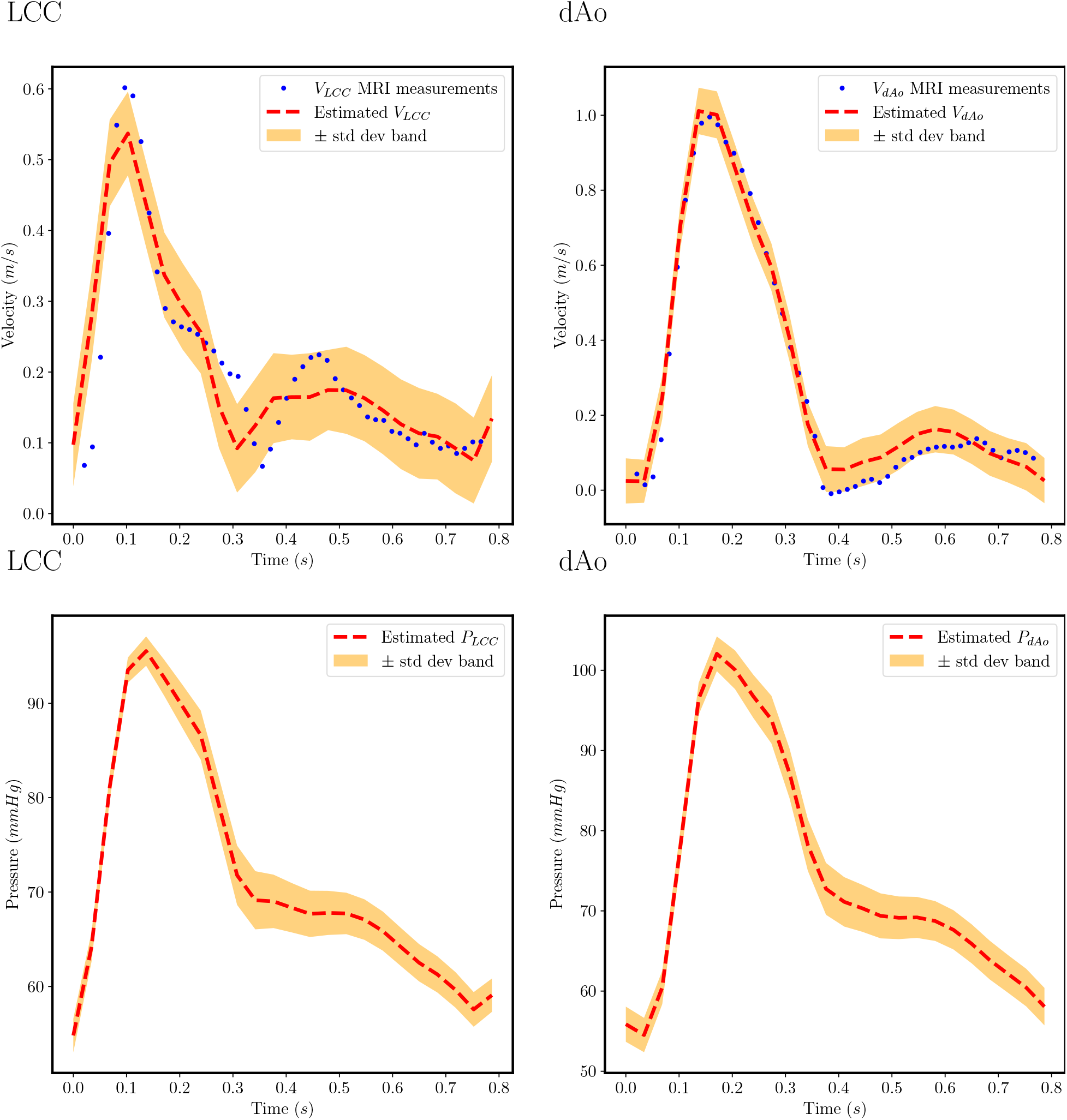
Resulting velocity and pressure wave-forms from aorta experiment using MRI flow data: In this figure the estimated wave-forms of the aorta geometry with MRI data is compared with the MRI data measurements. The MRI data are denoted by blue dots. The red line represents the waveform resulting from using the parameters that provide the maximum likelihood and the yellow area represents the model solution uncertainty.The abbreviation dAo correspond to the Descending Aorta and LCC to the Left Common Carotid arteries.

### 3.2 Maternal pelvic arterial results

We employ the computational model proposed in Section 2 for estimating the remodeling parameters in both the Normal subject and the pre-HPD subject cases. The predicted flow rate in the left and right external iliac arteries across 200 samples of the posterior distribution in the normal subject, are 950.18 ± 14.04 and 919.64 ± 38.92 mL/min, respectively. The predicted flow rate in the left and right uterine arteries are 323.02 ± 3.84 and 65.57 ± 0.98 mL/min, respectively (Figure 5). The pre-HPD subject has left and right external iliac artery flow rates 472.2 ± 16.5 and 711.02 ± 27.8 mL/min, respectively, and left and right uterine artery flow rate of 62.75 ± 4.5 and 134.22 ± 4.1 mL/min, respectively (Figure 7). When comparing between predicted and MRI-measured flow rates, the errors are highly variable (5.29-92.4% error). This discrepancy occurred because the model cannot provide an accurate prediction of the cross-sectional area of the right and left external iliac arteries. One reason for this lack of predictability is high errors are accumulated for the case of the cross-sectional area, as it is present in the denominator of the *β* parameter in the pressure equation of the model. Another reason is that we assume that the area measurement provided by the structural MRI is the equilibrium cross sectional area for the non-outlet vessels, left and right common iliac arteries and the descending aorta, which can contribute to inaccuracies. We present the predictions for the biomarkers computed from MRI, Bayesian inference and, also, include the Doppler Ultrasound derived biomarkers on Table 6. The predicted PI and RI values of the Normal subject are collectively lower than that of the pre-HPD subject. This trend agrees with the corresponding MRI and ultrasound-based PI and RI comparisons.

**Figure 5:**
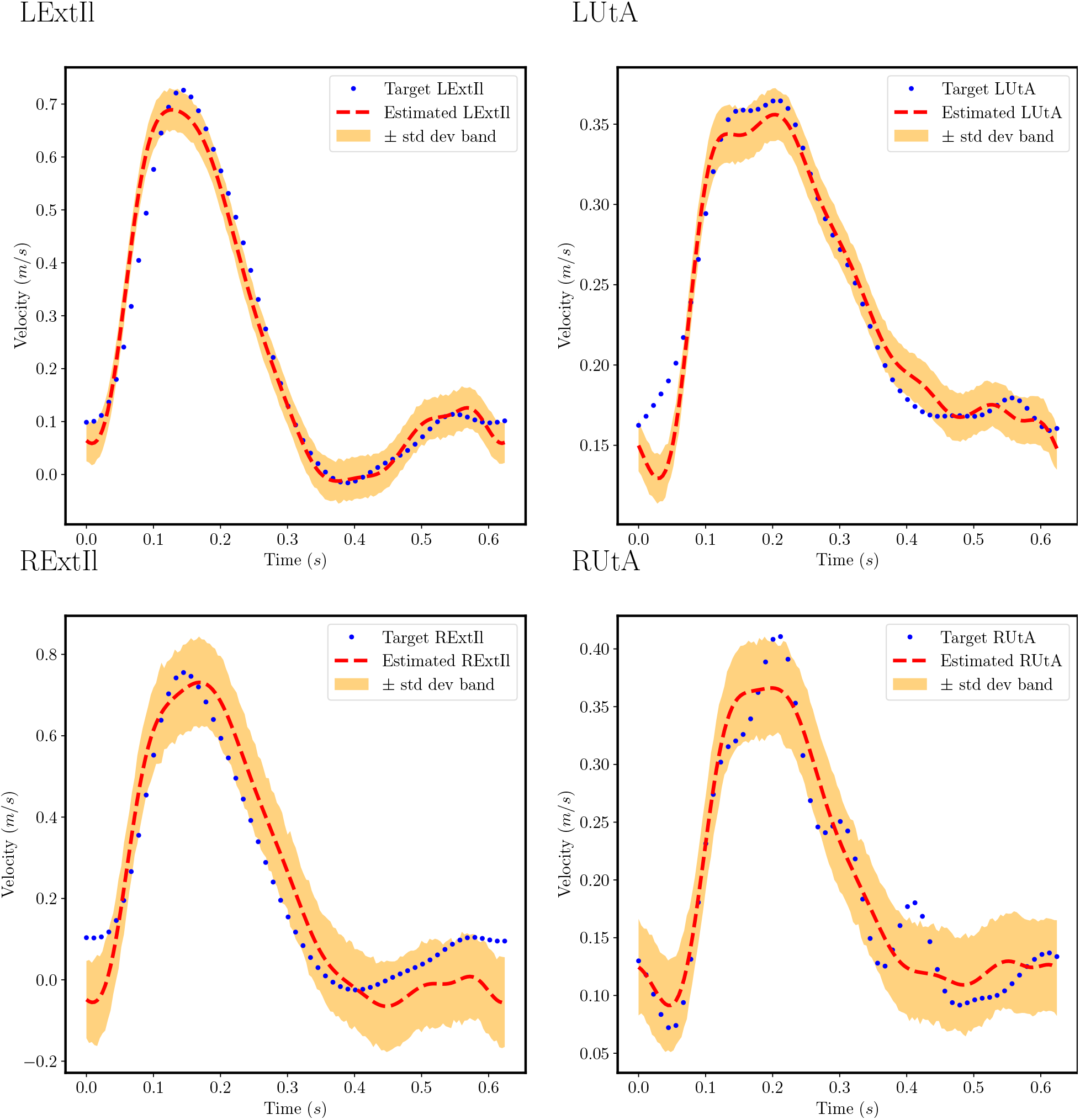
Resulting velocity wave-forms from maternal pelvic arterial experiment of Normal pregnant subject: In this figure the estimated wave-forms of the maternal pelvic arteries for Normal subject are compared with the interpolated mean velocity measurements from 4D flow MRI. The MRI measurements are denoted by blue dots. The red dashed lines represent the wave-forms resulting from using the parameters that provide the maximum likelihood. The abbreviation LExtIl correspond to the Left External Iliac, RExtIl correspond to the Right External Iliac, LUtA correspond to the Left Uterine and RUtA to the Right Uterine arteries.

The predicted R and C values are reported in Table 5. The predicted R values for the pre-HPD subject are higher than the R values of the Normal subject in the left external iliac artery, left uterine artery and right external iliac artery. The predicted compliance for bilateral external iliac arteries is lower in the pre-HPD subject compared to that of the Normal subject. However the opposite is true for bilateral uterine arteries, where the compliance is higher in the pre-HPD subject. Table 5 also reports the estimated cross-sectional area values. Both subjects have similar external iliac artery areas with the pre-HPD subject having a slightly narrower left external iliac artery. For the pre-HPD subject, both uterine artery cross-sectional areas are close to 25% of the external iliac artery areas. For the Normal subject we see a higher variability in the uterine cross-sectional areas with the left uterine artery to be higher. In terms of the predicted pressure wave-forms for both cases, they are (in general) higher for the pre-HPD subject case for systolic and diastolic pressures compared to the Normal subject case. In the left uterine artery, the predicted pressure is 152.4 ± 7.8/ 65.1 ± 7.3 mmHg vs. 107.78 ± 1.83/32.99 ± 1.49 mmHg, and in the right uterine artery, the predicted pressure is 148.2 ± 8.3 / 64.7 ± 7.14 mmHg vs. 85.78 ± 3.33 /31.0 ± 1.84 mmHg (see Figures 6 and 8). Moreover, we present comparative figures for the velocities and the pressures for all vessels for both the Normal and the pre-HPD subjects in figures 9 and 10 respectively. We observe that the velocities are collectively higher in the case of the Normal subject while the pressures are collectively higher in the case of the pre-HPD subject.

**Table 5:** Discovered resistance, compliance, and cross-sectional area values for both Normal and pre-HPD subjects: The estimated total arterial resistances (*R*), total arterial compliances (*C*) and equilibrium cross-sectional areas (*A*) from the Bayesian inference algorithm performed on the maternal pelvic arteries using mean velocities from 4D flow MRI are reported as mean ± standard deviation. The abbreviation LExtIl correspond to the Left External Iliac, RExtIl correspond to the Right External Iliac, LUtA correspond to the Left Uterine and RUtA to the Right Uterine arteries.

| Vessel Name | Subject | Normal subject |  |  | pre-HPD subject |  |  |
| --- | --- | --- | --- | --- | --- | --- | --- |
| | | $R \times 10^8 Pa s m^{-3}$ | $C \times 10^{-11} Pa^{-1} m^3$ | $A \times 10^{-6} m^2$ | $R \times 10^8 Pa s m^{-3}$ | $C \times 10^{-11} Pa^{-1} m^3$ | $A \times 10^{-6} m^2$ |
| LExtIl | | 4.64 ± 0.02 | 97.20 ± 11.95 | 68.11 ± 1.18 | 16.32 ± 3.39 | 35.79 ± 5.50 | 59.98 ± 7.15 |
| LUtA | | 14.23 ± 3.71 | 0.30 ± 0.17 | 22.68 ± 3.58 | 112.52 ± 14.72 | 3.2 ± 0.44 | 12.18 ± 1.41 |
| RExtIl | | 4.70 ± 0.19 | 96.26 ± 6.14 | 69.21 ± 0.70 | 7.38 ± 0.66 | 42.16 ± 2.71 | 68.87 ± 1.13 |
| RUtA | | 62.22 ± 2.38 | 2.76 ± 0.347 | 5.60 ± 0.32 | 43.76 ± 10.20 | 3.67 ± 0.53 | 16.97 ± 2.70 |

**Table 6:**
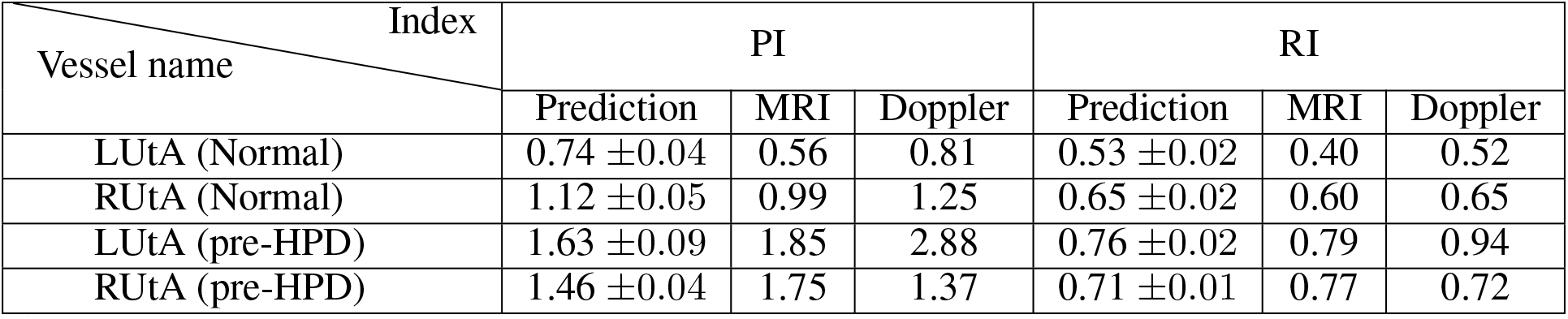
Comparison of the Pulsatility and Resistivity indices in Normal and pre-HPD pregnancies acquired from three different sources: In this table, we present the Pulsatility and Resistivity indices (PI, RI) of the left and right uterine arteries (LUtA, RUtA) acquired from MRI, model prediction, and Doppler ultrasound for both subjects. For the model predictions we present the mean values together with the standard deviations computed from 200 sampled velocity wave-forms from the posterior distribution. The abbreviation LExtIl correspond to the Left External Iliac, RExtIl correspond to the Right External Iliac, LUtA correspond to the Left Uterine and RUtA to the Right Uterine arteries.

| Vessel name \ Index | PI |  |  | RI |  |  |
| --- | --- | --- | --- | --- | --- | --- |
|  | Prediction | MRI | Doppler | Prediction | MRI | Doppler |
| LUtA (Normal) | $0.74 \pm 0.04$ | 0.56 | 0.81 | $0.53 \pm 0.02$ | 0.40 | 0.52 |
| RUtA (Normal) | $1.12 \pm 0.05$ | 0.99 | 1.25 | $0.65 \pm 0.02$ | 0.60 | 0.65 |
| LUtA (pre-HPD) | $1.63 \pm 0.09$ | 1.85 | 2.88 | $0.76 \pm 0.02$ | 0.79 | 0.94 |
| RUtA (pre-HPD) | $1.46 \pm 0.04$ | 1.75 | 1.37 | $0.71 \pm 0.01$ | 0.77 | 0.72 |

**Figure 6:**
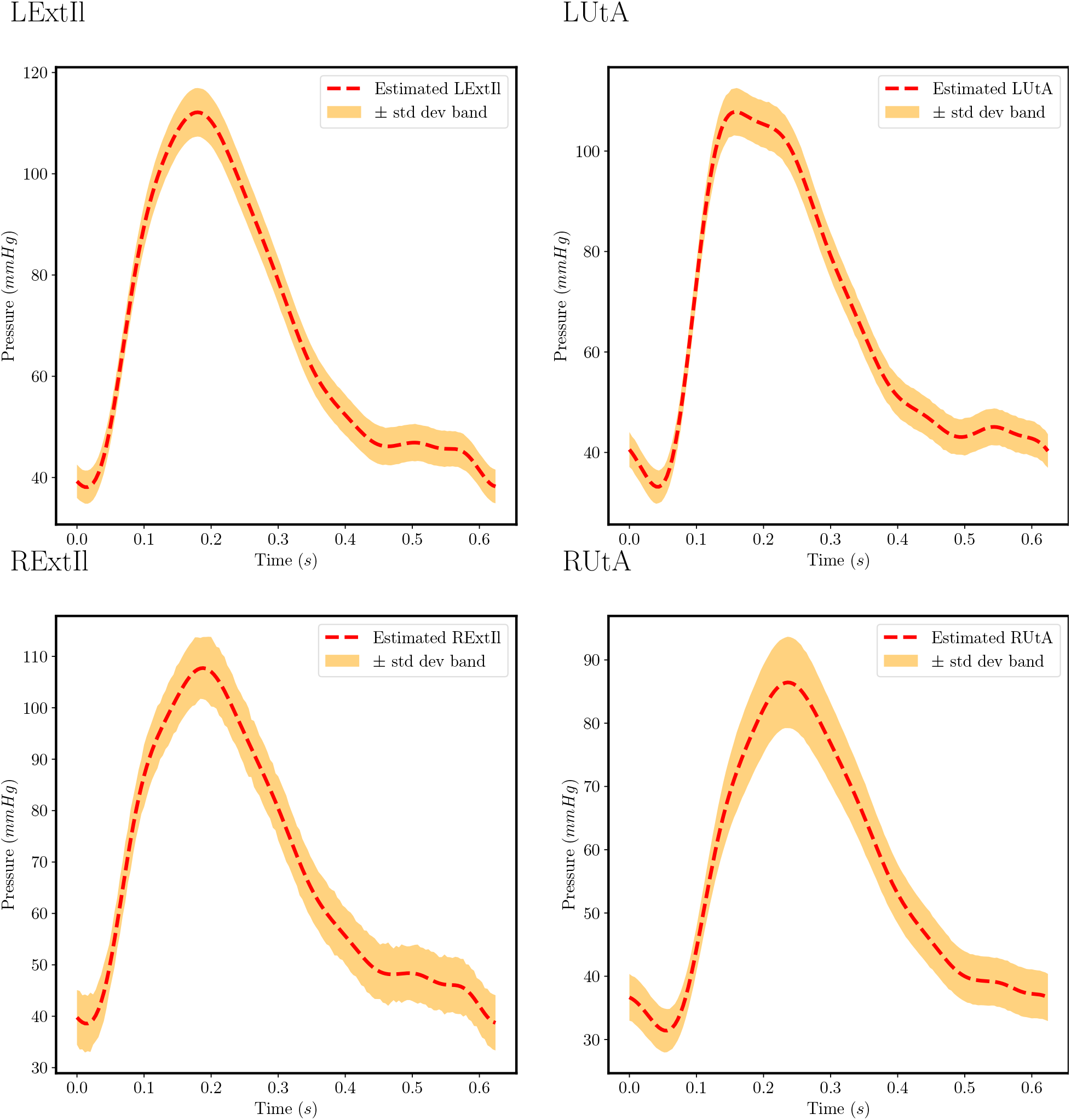
Resulting pressure wave-forms from maternal pelvic arterial experiment of Normal pregnant subject: In this figure the estimated pressure wave-forms of the maternal pelvic arteries for Normal subject are presented. The red dashed lines represent the wave-forms that result from using the parameters that provide the maximum likelihood and the yellow shaded areas are the model uncertainty. The abbreviation LExtIl correspond to the Left External Iliac, RExtIl correspond to the Right External Iliac, LUtA correspond to the Left Uterine and RUtA to the Right Uterine arteries.

**Figure 7:**
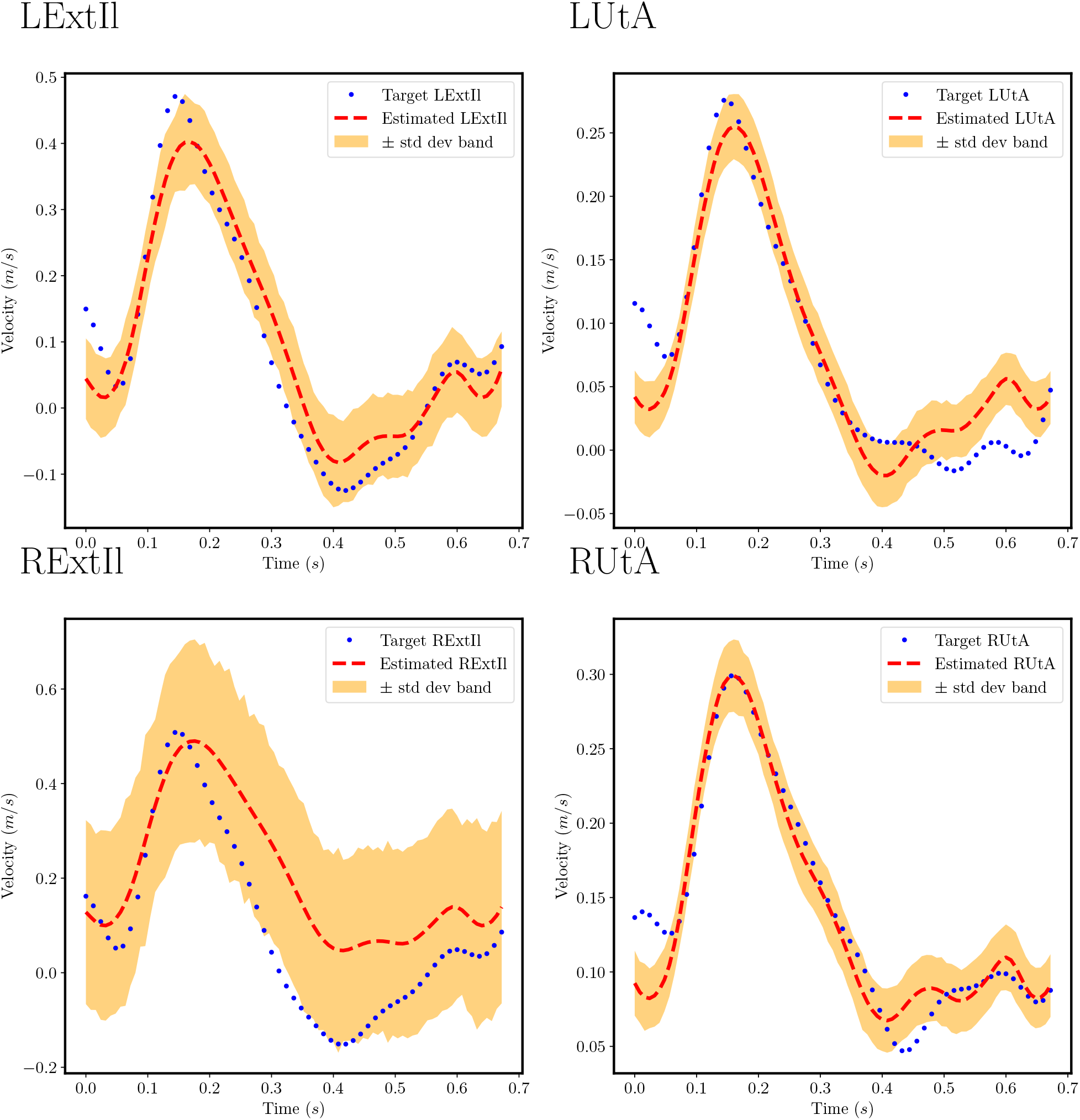
Resulting velocity wave-forms from maternal pelvic arterial experiment of pre-HPD subject: In this figure the estimated wave-forms of the maternal pelvic arteries for pre-HPD subject are compared with the interpolated mean velocity measurements from 4D flow MRI. The MRI measurements are denoted by blue dots and the red dashed lines represent the velocity wave-forms resulting from the parameters that provide the maximum likelihood and the yellow shaded areas are the model uncertainty. The abbreviation LExtIl correspond to the Left External Iliac, RExtIl correspond to the Right External Iliac, LUtA correspond to the Left Uterine and RUtA to the Right Uterine arteries.

**Figure 8:**
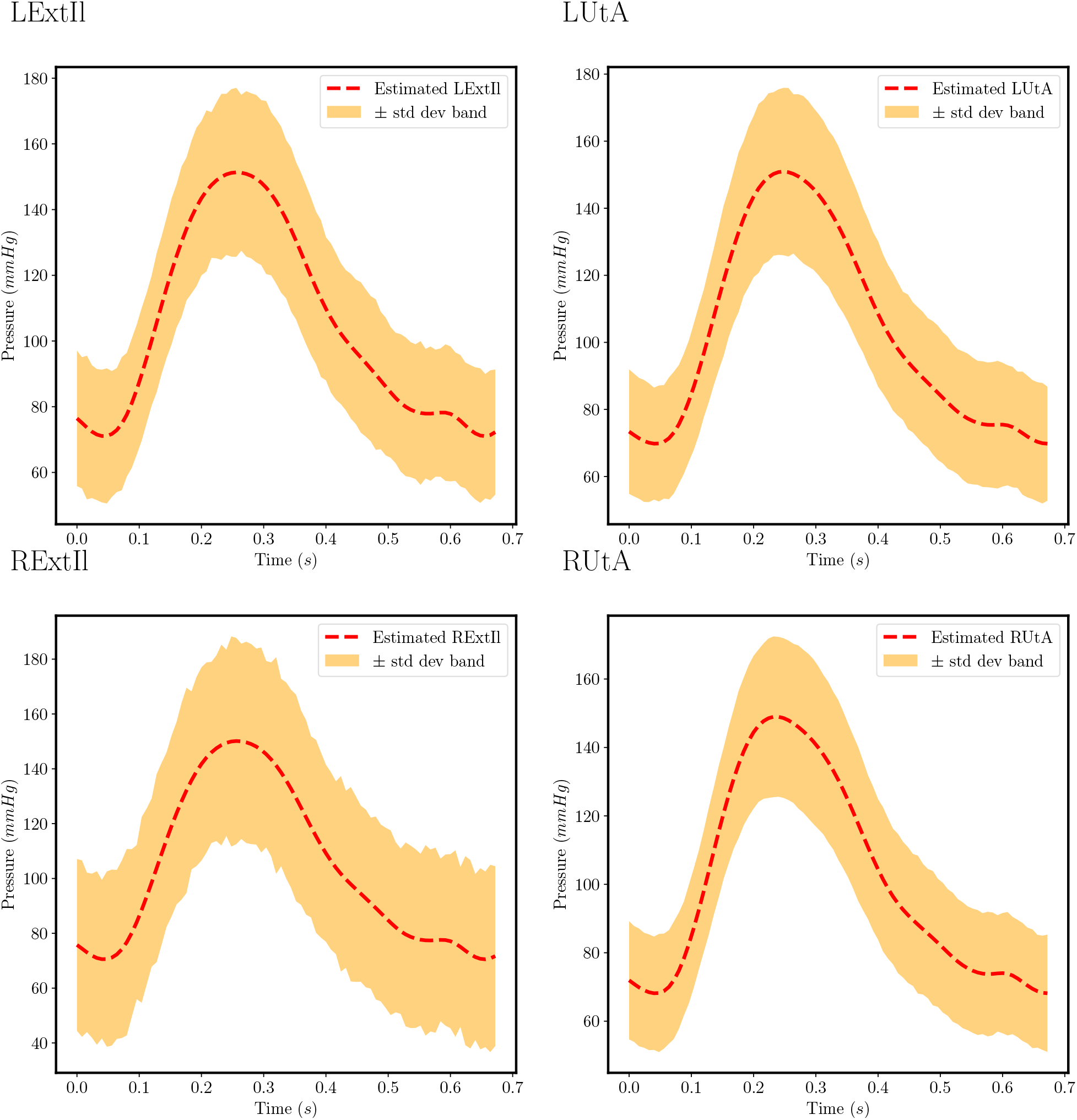
Resulting pressure wave-forms from maternal pelvic arterial experiment of pre-HPD subject: In this figure the estimated pressure wave-forms of the maternal pelvic arteries for pre-HPD subject are presented. The red dashed lines represent the wave-forms that result from using the parameters that provide the maximum likelihood value and the yellow shaded areas are the model uncertainty. The abbreviation LExtIl correspond to the Left External Iliac, RExtIl correspond to the Right External Iliac, LUtA correspond to the Left Uterine and RUtA to the Right Uterine arteries.

**Figure 9:**
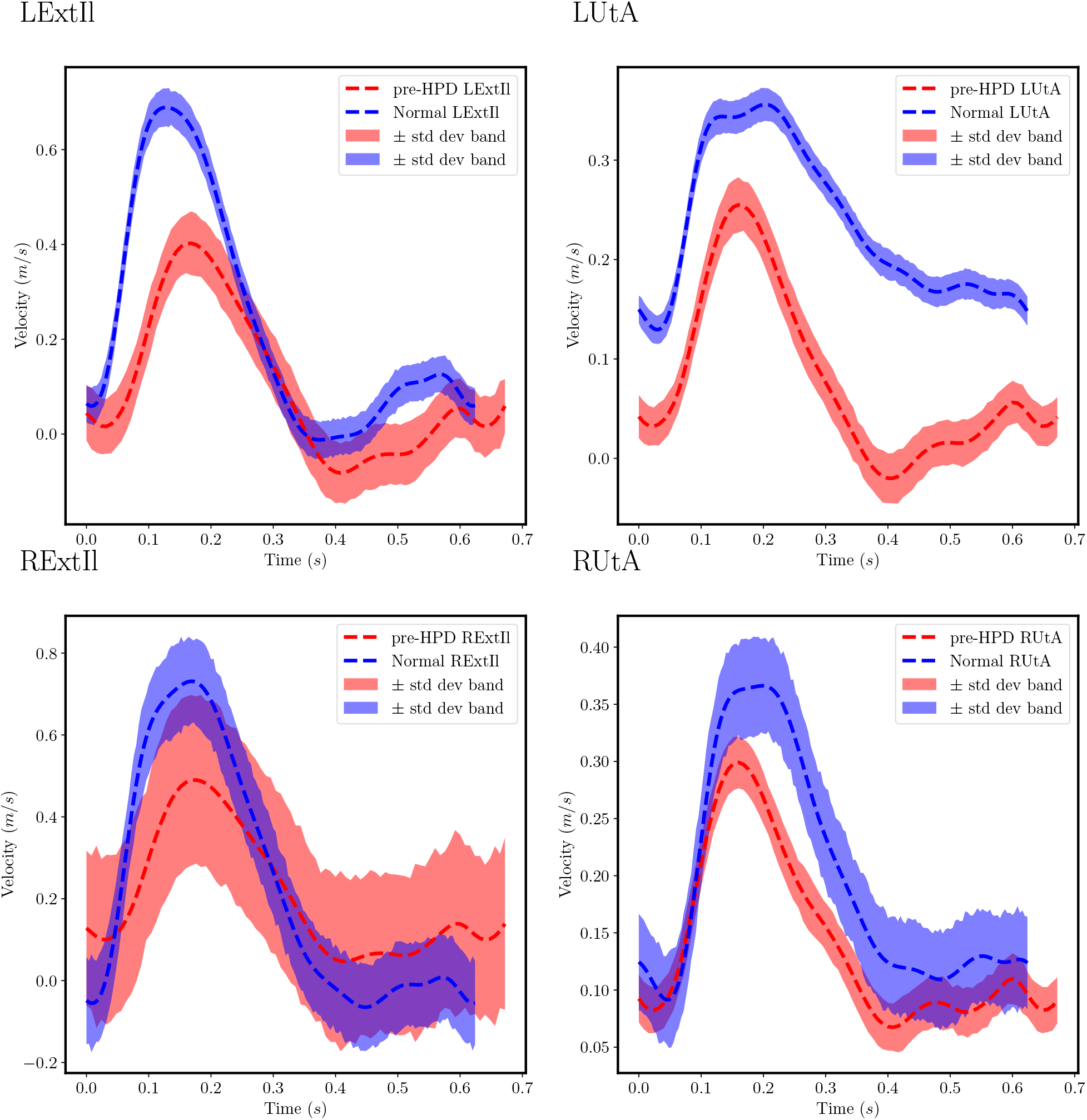
Comparison between predicted velocity wave-forms from maternal pelvic arterial experiment of Normal and pre-HPD pregnant subjects: In this figure the estimated wave-forms of the maternal pelvic arteries for both the Normal and the pre-HPD subjects are compared. The red and blue dashed lines represent the wave-forms resulting from using the parameters that provide the maximum likelihood for the pre-HPD and Normal subjects respectively. The abbreviation LExtIl correspond to the Left External Iliac, RExtIl correspond to the Right External Iliac, LUtA correspond to the Left Uterine and RUtA to the Right Uterine arteries.

**Figure 10:**
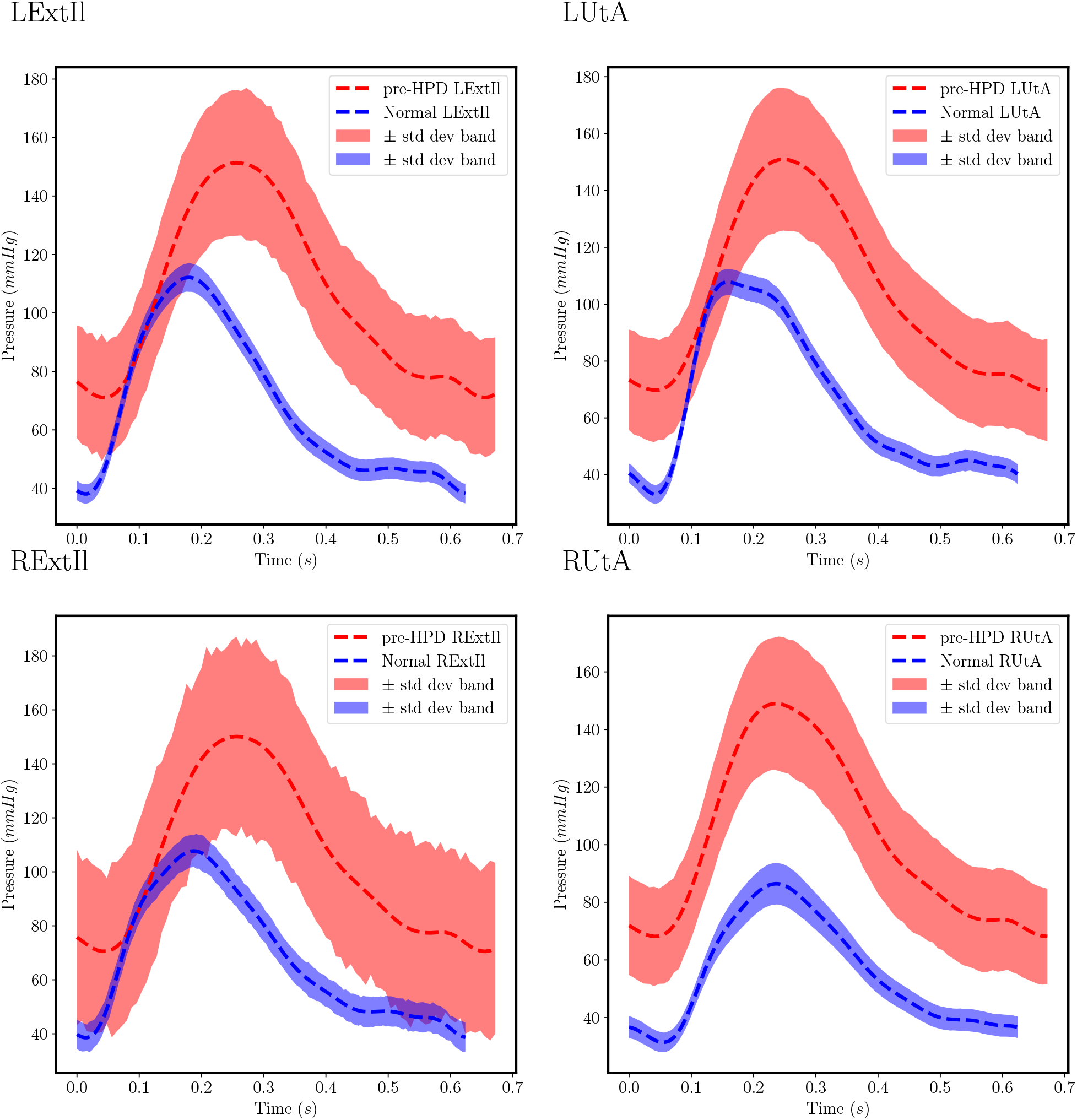
Comparison between predicted pressure wave-forms from maternal pelvic arterial experiment of Normal and pre-HPD pregnant subjects: In this figure the estimated wave-forms of the maternal pelvic arteries for both the Normal and the pre-HPD subjects are compared. The red and blue dashed lines represent the wave-forms resulting from using the parameters that provide the maximum likelihood for the pre-HPD and Normal subjects respectively. The abbreviation LExtIl correspond to the Left External Iliac, RExtIl correspond to the Right External Iliac, LUtA correspond to the Left Uterine and RUtA to the Right Uterine arteries.

## 4 Discussion

This study demonstrates the feasibility of inferring remodeling parameters of the maternal pelvic arterial network from MRI geometry and velocity data within a computational blood flow modeling framework. By adopting a Bayesian inference approach, we are able to iteratively match the predicted velocity wave-forms with the measured velocity wave-forms in a favorable manner. By doing so, we are able to recover parameters such as resistance, compliance, and pressure, which are not measured but inferred from physical principles. We initially perform proof-of-concept experiments in an aorta and carotid data set of a healthy volunteer and show that the Bayesian inference algorithm matched the target synthetic flow wave-forms and the target MRI flow wave-forms. There is greater uncertainty in the predicted velocity wave-forms for the aorta model with MRI flow data potentially because of the measurement noise present in the dataset. In the synthetic data case, the target resistance and compliance are chosen to be the same as the ones computed by a different estimation technique proposed by Kissas *et. al.*^30^, when applied to the real MRI descending aorta case. We choose these values because we are already aware that the results are in physiological ranges. Both the target and the predicted parameters are similarly matched in both outlets. In both aorta models, the left common carotid artery had lower predicted resistance and higher compliance than the descending aorta. The predicted pressure wave-forms of the descending aorta and left common carotid artery are also within physiological range based on cardiac catheterization measurements in a human^46^.

In the maternal pelvic arterial models, the Bayesian inference algorithm successfully recovered the remodeling parameters despite greater complexity in the network. Traditional measures of PI and RI show that the subject who went on to develop preeclampsia (pre-HPD subject) has elevated second trimester uterine artery pulsatility compared to the normal case (Normal subject), a trend which is often interpreted as higher resistance. When comparing the predicted resistance values between the models for the Normal subject and the pre-HPD subject, the pre-HPD subject has higher resistance than the Normal subject in the left uterine, the right and the left external iliac arteries, which indicates potential incomplete remodeling of these vessels in this case. On the other hand, the right uterine artery has higher resistance in the Normal subject potentially because it has not remodeled yet. Previous studies have described reduced vascular tone as part of the general pattern of arterial remodeling in pregnancy^10^. Since preeclampsia has been described to involve insufficient conversion of the spiral arteries from pre-pregnant vasoactive vessels to flaccid conduits^5^, we postulated that HPD results from failure to also completely lower the compliance of the uterine arteries. However, our results showed the pre-HPD subject case had higher compliance than the Normal subject case for the uterine arteries. Although the discrepancy between our results and this theory can be attributed to patient variability and the pilot nature of this initial study, animal studies have indeed found higher myogenic tone in healthy pregnant sheep and guinea pigs compared to healthy non-pregnant animals^10^, which seems to show that there may be a complex relationship between resistance and compliance and clinical outcomes that need to be further investigated. Moreover, the uterine artery is potentially exposed to more local vascular mediators from the placenta/pregnancy, possibly explaining why we may see remodeling/compliance changes there that are different from what is seen in systemic circulation^47^.

Unlike the aorta, the outlet cross-sectional areas for the maternal pelvic arterial network were also considered as inferred parameters because the 4D flow MRI did not have sufficient spatial resolution to precisely measure the areas of the small uterine arteries. The pre-HPD subject has lower cross-sectional area in the left uterine artery compared to the Normal subject, while higher in the the right uterine artery. Given the hypothesis that smaller luminal areas correlated with poorly remodeled uterine arteries in preeclampsia^10,21^, it made sense that the pre-HPD subject would have low cross-sectional areas of the uterine arteries, whereas the Normal subject would have at least one dilated uterine artery. For the right uterine artery, the predicted cross-sectional area of the Normal subject is smaller than the pre-HPD subject, which might be an indication that the particular artery has yet to start remodeling. The pre-HPD subject had consistently higher predicted pressure in all four outlets compared to the Normal subject.

There are several potential reasons for the discrepancies observed between the predicted and target velocity wave-forms. First, the measured velocities from MRI have a significant amount of noise, so they do not necessarily obey the underlying physics of the Navier-Stokes equations. Second, we performed simulations on a single subject using a single measurement and therefore did not have the advantage of averaging velocities over a population which would have improved robustness. Third, we assumed that the curvature of the vasculature is small enough to be considered one dimensional. We also, assumed that the velocity magnitude in the direction of the flow is much larger than in any other directions. These assumptions were made in order to reduce the model from three-dimensional to one-dimensional. Due to the complexity of real flows, these assumptions do not hold completely, therefore we have lost accuracy and introduced model discrepancy in the process. Finally, the low spatio-temporal 4D flow MRI resolution in the region of the maternal pelvic arterial network introduced uncertainties due to the inaccuracies in structural parameter measurements, such as the vessel cross-sectional area, and the small number of velocity measurements comprising a cardiac cycle. These modeling assumptions and measurement errors might result to phenomena where the uncertainty provided for the model might be high for some parameter of interest, i.e. see the right external iliac artery prediction in figure 7. These shortcomings of the method, which are related with the imaging techniques and technology, could be potentially alleviated by the advances in medical imaging technologies, paving the way for more accurate parameter estimation and potentially the successful enhancement of the proposed methodology in higher dimensions (i.e. 3D flow data). Parameter inference with higher dimensional data is possible but not efficient at this point, one reason being that the lack of measurement accuracy lessens the convergence speed of the inference methods.

A limitation of this study was that the geometry of the maternal pelvic arterial models did not include all the branches of the internal iliac arteries aside from the uterine arteries. While it is known that a key feature of maternal physiological adaptation to pregnancy is diverting more blood flow to the uterine arteries^6^, this simplification may have caused inaccuracies without including the other branches. A more comprehensive model would require improvements in the spatial resolution of the 4D flow MRI technique to measure velocities in the other branches or at least inclusion of their structures so that the additional outlet velocities can be inferred. Another limitation is that this study reported total peripheral resistance, rather than separating between characteristic impedance and the systemic peripheral resistance. This separation can be useful in future studies to determine the source of the resistance, whether in the larger arteries of the pelvis or the small uteroplacental vessels (e.g. arcuate, radial, spiral arteries), which can affect interpretation of the remodeling changes based on computational results. It should also be noted that two large vessels, the brachiocephalic trunk and left subclavian artery, are missing from the aorta geometry, which might affect the results of this study. However, the fact that our estimated pressures were within physiologic range is reassuring regarding the validity and applicability of our chosen methods.

This study demonstrated the use of Bayesian inference to predict parameters that are not routinely measured clinically but can be helpful in identifying high risk pregnancies in early gestation with the aid of MR imaging and computational fluid dynamics. In future work, more patients are needed to investigate the relationship between pressure, resistance, compliance, cross-sectional area, and disease characteristics. This will aid in determining the most influential parameters that can be correlated with clinical outcomes. These parameters are promising biomarkers because they may be more physiologically relevant compared to PI and RI.

## 5 Conclusions

## 6 Acknowledgements

We are grateful to Brianna Moon and Veronica Aramendia Vidaurreta for their help in acquiring the aorta MRI data. The maternal pelvic arterial network MRI data were acquired with the help of MRI technologists, research coordinators, and pregnant research participants at the Hospital of University of Pennsylvania. G.K. and P.P. would like to acknowledge support from the US Department of Energy under the Advanced Scientific Computing Research program (grant DE-SC0019116) and Air Force Office of Scientific Research (grant FA9550-20-1-0060). E.H. would like to acknowledge support from the National Institute of Health (grant 1F31HD100171). E.H., N.S., W.W. and J.D. acknowledge support from National Institute of Health (grant U01HD087180). E.W.T acknowledges support from NIH Medical Scientist Training Program T32 GM07170.

## Notes

### Competing Interest Statement

The authors have declared no competing interest.

